# Differential yet integral contributions of Nrf1 and Nrf2 in the human antioxidant cytoprotective response against *tert*-butylhydroquinone as a pro-oxidative stressor

**DOI:** 10.1101/2021.06.05.447190

**Authors:** Reziyamu Wufur, Zhuo Fan, Keli Liu, Yiguo Zhang

## Abstract

In the past 25 years, Nrf2 had been preferentially parsed as a master hub of regulating antioxidant, detoxification and cytoprotective genes, albeit as a matter of fact that Nrf1, rather than Nrf2, is indispensable for cell homeostasis and organ integrity during normal growth and development. Here, distinct genotypic cell lines (*Nrf1α^−/−^, Nrf2^−/−ΔTA^* and *caNrf2^ΔN^*) are employed to determine differential yet integral roles of Nrf1 and Nrf2 in mediating antioxidant responsive genes to *t*BHQ as a pro-oxidative stressor. In *Nrf1α^−/−^* cells, Nrf2 was highly accumulated but also cannot fully compensate specific loss of Nrf1α’s function in its basal cytoprotective response against endogenous oxidative stress, though it exerted partially inducible antioxidant response, as the hormetic effect of *t*BHQ, against apoptotic damages. By contrast, *Nrf2^−/−ΔTA^* cells gave rise to a substantial reduction of Nrf1 in both basal and *t*BHQ-stimulated expression and hence resulted in obvious oxidative stress, but can still be allowed to mediate a potent antioxidant response, as accompanied by a significantly decreased ratio of GSSG to GSH. Conversely, a remarkable increase of Nrf1 expression was resulted from the constitutive active *caNrf2^ΔN^* cells, which were not manifested with oxidative stress, no matter if it was intervened with *t*BHQ. Such inter-regulatory effects of Nrf1 and Nrf2 on antioxidant and detoxification genes (encoding HO-1, NQO1, GCLC, GCLM, GSR, GPX1, TALDO, MT1E and MT2), as well on the ROS-scavenging activities of SOD and CAT, were further investigated. The collective results unraveled that Nrf1 and Nrf2 make distinctive yet cooperative contributions to finely tuning basal constitutive and/or *t*BHQ-inducible expression levels of antioxidant cytoprotective genes in the inter-regulatory networks. Overall, Nrf1 acts as a brake control for Nrf2’s functionality to be confined within a certain extent, whilst its transcription is regulated by Nrf2.

## 1. Introduction

With the development of science and technology, much more effective compounds with antioxidant properties have been discovered or synthesized insofar as to prevent lipid and protein oxidation [1, 2]. Amongst them, *tert*-butylhydroquinone (*t*BHQ) is a known small molecule phenolic antioxidant, which is a main metabolite of 3-*tert*-butyl-hydroxyanisole (BHA) in *vivo* in humans, dogs and rats, since it is widely used as a preservative in oils and processed foods [3, 4]. However, *t*BHQ (and its precursor BHA) is *de facto* identified as a double-faced compound with both effects to be exerted as an antioxidant and also a pro-oxidant in biological systems [5, 6]. Of note, such a double-bladed sword impact of *t*BHQ is further unraveled by chemoprotective and carcinogenic effects of this compound and its reactive metabolites [6], in addition to its cytotoxicity [3, 4]. This is due to the fact that oxidative metabolism of *t*BHQ by metal-mediated redox cycling and microsomal monooxygenase system [e.g. phase I drug-metabolic enzyme cytochrome P450 1a1 (Cyp1a1)] yields several reactive oxygen species (ROS) and electrophilic intermediates, followed by the formation of reactive glutathione (GSH)-conjugates [e.g. in phase II drug-metabolic reactions by glutathione-S transferases (GSTs)]. Furtherly, *t*-BHQ is also identified to function as a novel ligand of aryl hydrocarbon receptor (AhR)[7], such that it can directly induce the expression of Cyp1a1, an enzyme known to play an important role in the chemical activation of xenobiotics to carcinogenic derivatives. Thereby, it is inferred that the AhR-dependent induction of Cyp1a1 by *t*BHQ represents a positive feedback network so as to promote carcinogenicity, particularly upon its long-term exposure, in the gastrointestinal and liver tissues [6, 8].

On the another facet, the cytoprotective effect of *t*BHQ on biological systems is revealed by *bona fide* induction of the endogenous antioxidant and detoxification genes (e.g., those encoding phase II drug-metabolic enzymes) in response to this food additive[8, 9]. The endogenous antioxidant defense is provided predominantly by reduced glutathione (GSH) and other thiol-sensitive signaling molecules (e.g., thioredoxin), which contribute to metabolism of potentially harmful pro-oxidant agents (e.g., xenobiotics) and restore the intracellular redox balance to a steady-state, so that cell homeostasis is rebalanced [5, 10]. Of note, the expression of such innate antioxidant biosynthetic and detoxifying enzymes is governed primarily by the Cap’n’Collar (CNC) basic region-leucine zipper (bZIP) family of transcription factors [11–13]. Amongst this family, Nrf1 and Nrf2 (both also called Nfe2l1 and Nfe2l2, respectively) are two principal regulators for maintaining robust redox homeostasis in life process [13–16]. To date, most studies of *t*BHQ-induced antioxidant cytoprotective responses are focused disproportionately on the redox-sensitive Nrf2 [8, 11, 17–19], rather than putative redox threshold-setting Nrf1 [20–22], since the former Nrf2 was firstly identified as a master regulator of the phase II detoxifying enzyme genes (e.g. *NQO1, HO-1, GSTs*) through their antioxidant response elements (AREs) to the pro-oxidant BHA or its metabolite *t*BHQ [5, 23]. The underlying mechanisms for *t*BHQ-stimulated activity of Nrf2 are well documented [11, 24, 25], but it is less understood whether and/or how the transactivation activity of Nrf1 is induced by exposure to this chemical.

Although Nrf2 is accepted as a master regulator of ARE-driven cytoprotective gene expression [11, 25], it is not essential for normal development and healthy growth, because its global knockout (*Nrf2^−/−^*) mice are manifested with neither any obvious defects nor spontaneous pathological phenotypes (e.g. cancer) [23, 26]. In effect, *Nrf2^−/−^* mice are more susceptible than wild-type mice to chemical carcinogens [27], in addition to oxidative stress [28]. Thereafter, induction of Nrf2 (by *t*BHQ) has been recognized as a potential chemopreventive and therapeutic target against cancer [25, 29]. To the contrary, the long-term induction of hyperactive Nrf2 is also reconsidered as a potent oncogenic driver with the hallmarks of cancer; this is based on its *bona fide* tumor-promoting effects and resistance to chemotherapy [30, 31]. Such dual opposing roles of Nrf2 in cancer prevention and progression should be taken into account for its bidirectional potentials to implicate in cancer treatment.

By sharp contrast, Nrf1 is endowed with its innate unique features that are distinctive from Nrf2 [12, 32, 33], as evidenced by its gene-targeting knockout (*Nrf1^−/−^*) to establish distinct animal models with significant pathological phenotypes [16, 20, 22, 34–36]. Global knockout of *Nrf1^−/−^* leads to embryonic lethality at E6.5 to E14.5, resulting from severe oxidative stress [20, 34, 35]. This fact implies that loss of Nrf1 cannot be compensated by Nrf2, though Nrf2 can also contribute to combinational regulation of antioxidant cytoprotective genes as confirmed by a double knockout model of *Nrf1^−/−^:Nrf2^−/−^* [14]. Furtherly, distinct tissue-specific *Nrf1^−/−^* mice are manifested with typical pathologies, resembling human non-alcoholic steatohepatitis (NASH) and hepatoma [16, 22], type-2 diabetes [37] and neurodegenerative diseases [38, 39]. Collectively, these demonstrate that mouse Nrf1 (and its isoforms) fulfills an indispensable function in regulating critical genes for maintaining robust redox homeostasis and organ integrity, so that the normal physiological development and growth are perpetuated in life process. However, it is regretable that these achievements are made mostly from mouse models. Such being the case, the underlying mechanism(s) by which human Nrf1 (or its derived isoforms) contributes to similar pathophysiological cytoprotective responses remains elusive.

For this reason, we have established three specific-knockout cell lines by gene-editing of human *Nrf1* or *Nrf2* on the base of HepG2 cells (named *Nrf1α^−/−^, Nrf2^−/−ΔTA^* and *caNrf2^ΔN^*, respectively) [31, 40]. Herein, these three distinct genotypic cell lines together with wild-type cells were stimulated by *t*BHQ and subjected to a series of experimental interrogation of both basal and inducible expression levels of certain antioxidant, detoxification and cytoprotective genes. The resulting evidence has been presented by us, revealing that human Nrf1 and Nrf2 make differential, yet integral, contributions to synergistic regulation of antioxidant and detoxification genes induced by *t*BHQ as a pro-oxidative stressor. Of great note, it is plausible that the presence of Nrf1 determines basal redox steady-state and normal antioxidant cytoprotective responses against endogenous oxidative damages and apoptosis, albeit Nrf2 is involved in this homeostatic function. This study also provides a better understanding of the inter-regulatory roles of Nrf1 and Nrf2 within the redox control system, except that both factors can exert their specific yet combinational functions in the process. Such cautions should be severely taken into account for us to develop new drugs targeting either Nrf1 or Nrf2 alone or both in the future biomedical study.

## 2. Materials and Methods

### 2.1 Cell lines and Regents

The human hepatomacellular carcinoma (HepG2) cells (i.e. *Nrf1/2^+/+^, WT*) were purchased from American Type Culture Collection (ATCC, Manssas, VA, USA). And three HepG2-derived cell lines with distinct knockout types of *Nrf1α^−/−^* (with a specific deletion mutant of full-length Nrf1/TCF11 and its derived isoforms), *Nrf2^−/−ΔTA^* (lacking its longer transactivation domain-containing fragment), *or caNrf2^ΔN^* (i.e. a constitutive active mutant of Nrf2 lacks its N-terminal Keap1-binding Neh2 domain) had been established in our laboratory, as described in detail by Qiu et al [31]. It is worth mentioning that the authenticity of HepG2 cell line had been confirmed by its authentication analysis and STR typing map (which was carried out by Shanghai Biowing Applied Biotechnology Co., Ltd., Shanghai, China). These cell lines were cultured in DMEM supplemented with 10% (*v*/*v*) FBS and 100 Units/L double-antibiotic (penicillin and streptomycin) and incubated at 37°C in a humidified 5% CO_2_ atmosphere.

Subsequently, those experimental cells were treated with *t*BHQ (CAS No.1948-33-0, from Sigma), which is a newly synthesized phenolic antioxidant with the chemical formula of *C_10_H_14_O_2_* at *MW* 166.22. This compound has completely dissolved in DMSO at a final concentration of 50 μM and stored at −20°C, before it is experimented. Of note, specific antibody against Nrf1 was made in our laboratory [41]. Besides, other five distinct antibodies against Nrf2 (ab62352), GCLC (ab207777), GCLM (ab126704), HO-1 (ab52947) or GPX1 (ab108427) were obtained from Abcam (Cambridge, UK). Additional antibodies against NQO1 (D26104), GSR (D220726) or TALDO1 (D623398) were from Sangon Biotech (Shanghai, China), whilst β-actin antibody (TA-09) was from ZSGB-BIO (Beijing, China).

### 2.2 Cell viability with MTT assay

All indicated experimental cells were digested by trypsin and diluted into a suspension of 5×10^4^ cell/mL, before being seeded into 96-well plates. After the cells were completely adherent to the plates, they were treated with *t*BHQ at different concentrations (i.e. 0-100 μM) for 24 h, or with 50 μM of *t*BHQ for distinct time periods (i.e. 0-24 h). The cell viability was evaluated by assaying 3-(4,5-dimethylthiazol-2-yl)-2,5-diphenyltetrazolium bromide (MTT, ST1537, Beyotime, Shanghai, China) to form an insoluble product formazan in living cells. The resulting data were calculated by dividing the experimental absorbance by relevant control values. The results were shown as a percentage of *mean ± SD* (*n=5*), which are representative of at least three independent experiments.

### 2.3 Quantitative RT-PCR analysis of mRNA expression

Experimental cells growing in logarithmic phase were digested with trypsin and diluted by a complete medium into the suspension of 3.5×10^5^ cell/mL. Then equal amounts of cells were inoculated in 6-well plates and cultured until being completely adherent. Thereafter, they were treated for different time periods (i.e. 0, 4, or 6 h) with 50 μM of *t*BHQ. Subsequently, total RNA was isolated by using a RNA extrection kit (TIANGEN, Beijing, China), 500 ng of which was then subjected to the reaction with reverse transcriptase (Promega, Madison, WI, USA) to synthesize the single strand cDNAs, that served as PCR templates. Basal and *t*BHQ-induced mRNA expression levels of those indicated genes were detected by quantitative real-time PCR (qRT-PCR) with each pairs of their primers (Table 1).

**Table 1.**
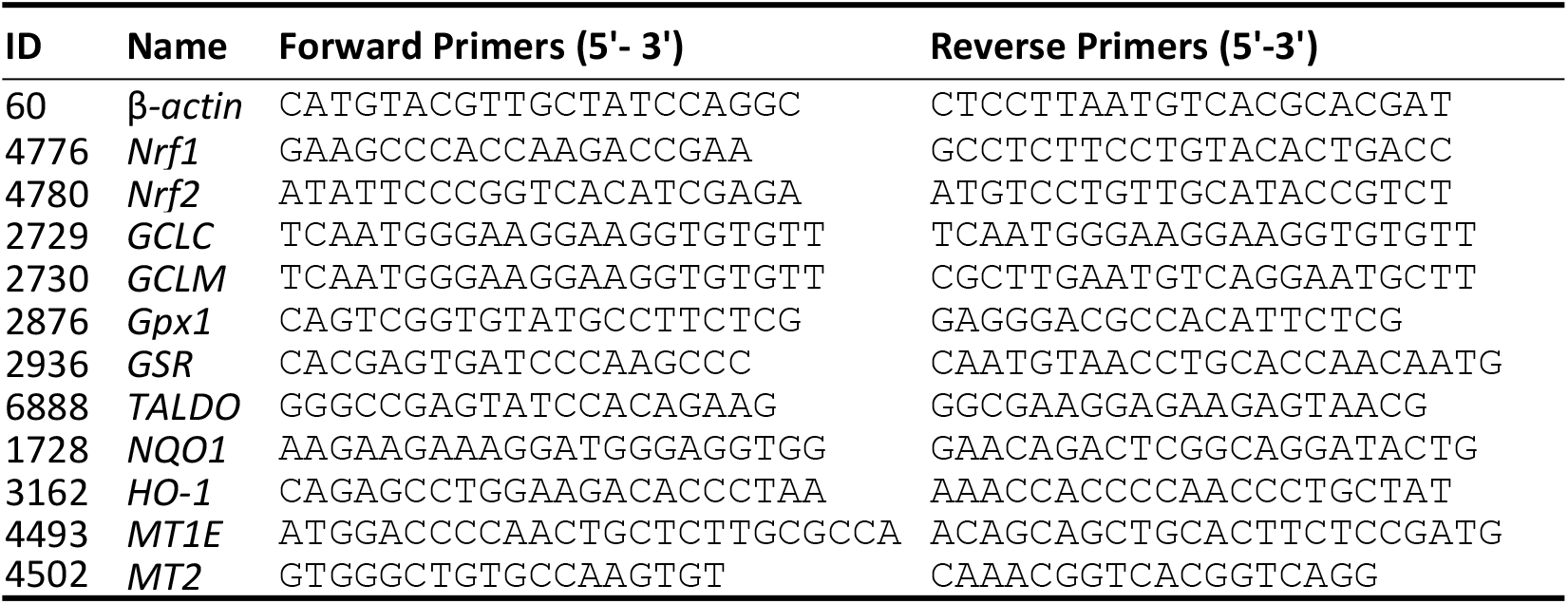
The primer pairs used for qRT-PCR analysis.

The qRT-PCR reaction was carried out with GoTaq^®^qPCR Master Mix (Promega, Madison, WI, USA) on a CFX96 instrument (Bio-Rad, Hercules, CA, USA). The specific reaction procedure was followed by all samples, which were inactivated at 95 °C or 3 min, and amplified by 40 reaction cycles of 15 s at 95°C and 30 s at 60 °C. The resulting data were analyzed by the Bio-Rad CFX 96 Manager 3.0 software (Hercules, CA, USA), whilst β-actin expression level served an internal reference control and experimental values obtained from treatment with *t*-BHQ for 0 h (i.e. *T*_0_) was used as relevant normal controls. The results of all examined genes were shown as fold changes (*Mean* ± *SD, n*=3), which are representative of at least three independent experiments.

### 2.4 Western blotting analysis of protein expression

After experimental cells were collected as described above, the proteins were extracted for total cell lysates, diluted with a 3× loading buffer and denatured by boiling at 100°C for 10 m. The samples from each of groups were electrophoretically separated by either 8% or 10% SDS-PAGE gels (running at 50 v for 30 min and then 100 v for 2 h), before being transformed on the PVDF membrane (at 200 A for 2 h). After the protein-botted membranes were blocked by 5% skimmed milk for 1 h, they were incubated with each of the indicated primary antibodies at 4 °C overnight, and then re-incubated with the secondary antibody at room temperature for 2 h. The protein blots were developed by the enhanced chemiluminescence as described previously [31]. The intensity of blots was calculated by using the Quantity One software (Bio-Rad Laboratories) and also normalized to β-actin as a loading control.

### 2.5 Detection of cellular ROS and apoptosis by flow cytometry

To determine intracellular ROS levels, all the cell samples were exposed to 100 μM dichlofluorescein diacetate (DCFH-DA included in the detection kit, S0033S, Beyotime, Shanghai, China) for 30 min at the incubator (37°C, 5% CO_2_). After being washed twice with PBS, the samples were centrifuged and resuspended in a serum-free medium. Then, the resulting 2’7’-dichlofluorescein was detected at the excitation wavelength of 488 nm and the emission wavelength of 525 nm by a flow cytometry (FlowJo, Ashland, OR, USA). In addition, the final results were expressed by the DCFH fluorescence intensity in distinct cells. Furthermore, those samples were collected by centrifuging at 1000× g for 5 min, and stained with binding buffer containing of Annexin V-FITC and propidium iodide (PI) for 15 min. After the samples were washed twice to remove the excess staining reagent, they were subjected to detection of cell apoptosis by flow cytometry. The resulting data were shown by different fluorescence intensity in distinct states of cells.

### 2.6 The assays for total, reduced and oxidized glutathione levels

After experimental cells had been treated with *t*BHQ for indicated times (i.e. 0, 4 or 6 h), all cell samples were collected in PBS and then subjected to the measurement of total glutathione, reduced glutathione (GSH) and oxidized glutathione (GSSG) by using a Glutathione assay kit (A061-1, Nanjing Jiancheng, Nanjing, China) according to the manufacturer’s instruction. Of note, two standards of GSH and GSSG were also prepared in the same assays. The assay was designed by using an Ellman’s reagent (*5,5’-disulfidebis −2-nitrobenzoic acid*, DNTB), which reacts with GSH to form *2-nitro-5-thiobenzoic acid*, a yellow products with a absorbance at wavelength of 405 nm. In addition, the protein concentrations in samples were determined by the bicinchoninic acid assay (BCA, P1511, ApplyGene, Beijing, China) and used as an internal control for the normalization, along with relevant standard curves, in order to calculate amounts of total glutathione, GSSG and GSH by the formula provided by this manufacturer. The final resulting data are shown by a ratio of GSSG to GSH levels.

### 2.7 Assays for ROS-scavenging activities of superoxide dismutase and catalase

All groups of experimental cells had been treated with *t*BHQ for distinct lengths of times (0, 4 or 6 h) before being collected and subjected to assays for ROS-scavenging activities of superoxide dismutase (SOD), that were determined according to the instruction of enhanced SOD assay kit (A001-3, Nanjing Jiancheng, China). The another ROS-scavenging enzyme catalase (CAT) activity was also detected through the instructions of CAT kit (BC0205, Solarbio, Beijing, China).

### 2.8 ARE-Luciferase Reporter Assays

All experimental cells were seed into 12-well plates, after reaching 80% confluence, the cells were co-transfected using a lipofectamine 3000 mixture with each of *ARE*-driven luciferase plasmids (which were made by inserting each of the indicated ARE sequences into the pGL3-Promoter vector) or non-ARE reporter plasmids (as background control), together with an experiment construct for Nrf1, Nrf2 or empty pcDNA3.1 vector. In this test, the *Renilla* expression by pRL-TK plasmide served as an internal control for transfection efficiency. And the luciferase activity was measured by the dual-luciferase reporter system (Beyotime, Shanghai, China).

### 2.9 Statistical analysis

All the results are presented as *mean* ± *SD* (*n=3 or 5*). The comparison of the various experimental groups and their corresponding controls was carried out by one-way ANOVA, and analyzed by the post-hoc test with Fisher’s least significant difference (LSD). The differences in between distinct treatments were considered to be statistically significant at *p*< 0.05.

## 3. Results

### 3.1 Different effects of Nrf1 and Nrf2 on cell growth during tBHQ intervention

To gain an insight into endogenous *Nrf1*- and *Nrf2*-mediated antioxidant responses to *t*BHQ, we here confirmed that four distinct genotypes of cell lines are true (Fig. 1A), as described previously [31, 40]. Of note, human *Nrf1α* is manifested with four major isoforms, as identified by Xiang *et al* [41], of which its A and B isoforms represent the full-length glycoprotein and deglycoprotein of Nrf1, respectively, whilst its C and D isoforms denote two distinct lengths of *Nrf1* N-terminally-truncated isoforms. Specific knockout of *Nrf1α* (by its gene-editing to delete a very short segment adjoining its translational start codons) led to a complete loss of all four *Nrf1α*-derived isoforms in *Nrf1α^−/−^* cells, albeit with a retention of other two minor proteins Nrf1^ΔN^ and *Nrf1β* (Fig. 1A). By contrast, all four *Nrf1α*-derived isoforms A to D were also substantially diminished by *Nrf2^−/−ΔTA^*, but its B to D isoforms (with distinct potentials of its *trans*-activity) were significantly augmented by *caNrf2^ΔN^*. Intriguingly, *Nrf1β* abundances were also markedly suppressed by *Nrf2^−/−ΔTA^* or *caNrf2^ΔN^*. These imply that Nrf1 expression and processing may be monitored by *Nrf2*, besides itself. Conversely, only a major *Nrf2* isoform-A, but not isoforms its B or C, was incremented in *Nrf1α^−/−^* cells (Fig. 1A). However, all three isoforms A to C of Nrf2 were completely abolished by specific deletion of its transactivation Neh4-Neh5 domains (to yield an inactive *Nrf2^−/−ΔTA^*), but their disappearance seemed to be replaced by another three smaller isoforms with a faster electrophoretic mobility when compared with equivalents examined in *WT* cells. By contrast, three slightly shorter isoforms of *caNrf2^ΔN^* (closely to wild-type isoforms A to C, respectively) were retained and enhanced by this constitutive active mutant factor, because the N-terminal Keap1-binding Neh2 domain of Nrf2 was removed from its genomic locus. Collectively, these indicate that Nrf2 expression and processing may also be monitored by itself, as well by Nrf1, within an inter-regulatory feedback cycle.

**Figure 1.**
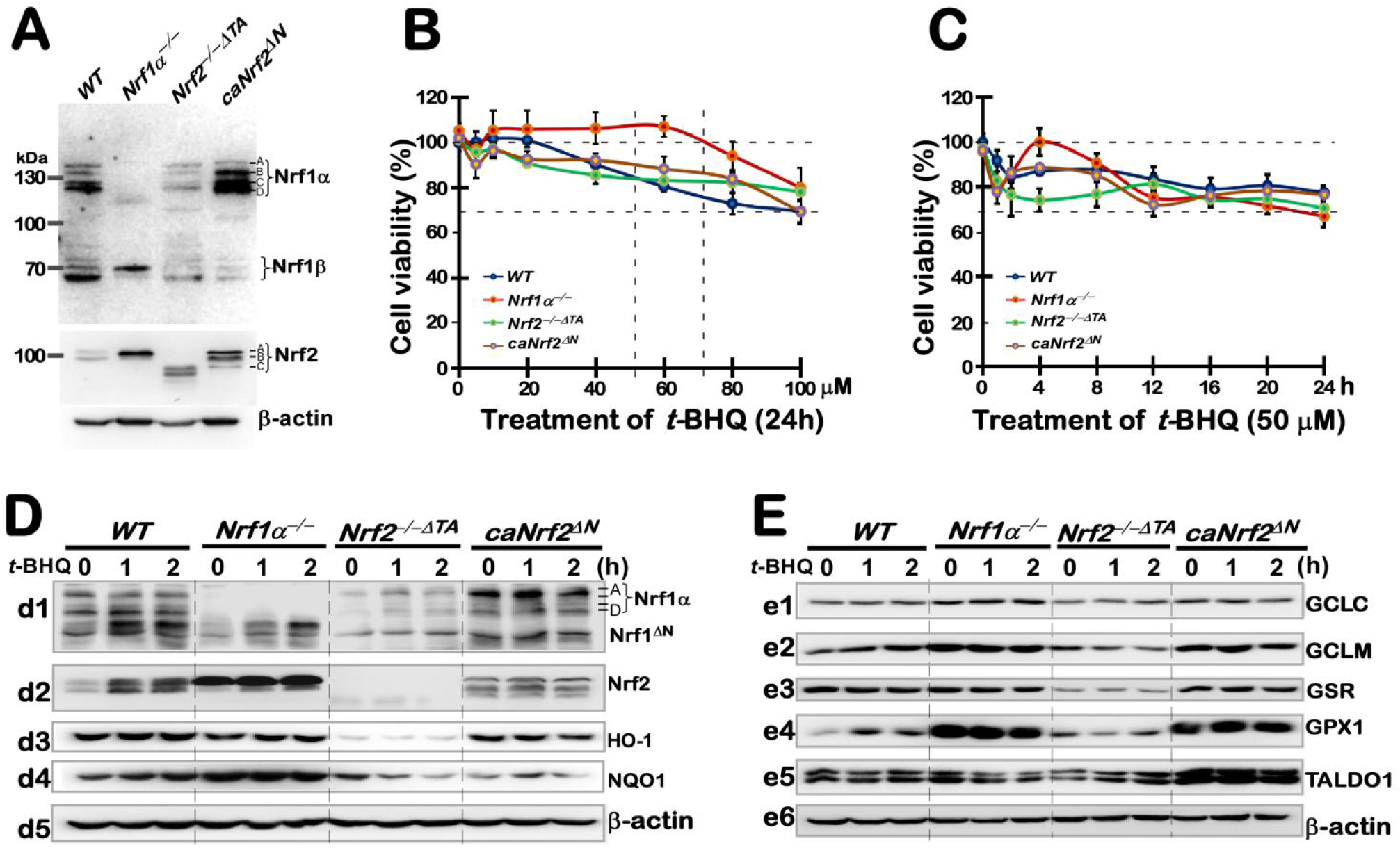
Distinct effects of *t*BHQ on different cell viability and antioxidant responsive genes. **(A)** Distinct protein abundances of Nrf1 and Nrf2 in *WT*, *Nrf1α^−/−^*, *Nrf2^−/−ΔTA^* or *caNrf2^ΔN^* cell lines were determined by western blotting with their specific antibodies. **(B** & **C)** Different cell viability was examined by the MTT-based assay, after the indicated cells had been treated with *t*BHQ: (*B*) at different doses (from 0 to 100 μM) for 24 h; (*C*) at a single dose of 50 μM for different length of time. **(D** & **E)** Four distinct cell lines were treated with 50 μM *t*BHQ or not for a short time (from 0 to 2 h), followed by western blotting of Nrf1, Nrf2 and other ARE-driven target gene expression levels as indicated.

Next, the cytotoxic effect of *t*BHQ on aforementioned four different cell lines was evaluated by a MTT assay for the formation of formazan precipitates with succinate dehydrogenase in the mitochondria of all living cells only, and changes in the absorbance were measured to reflect the cell viability. As shown in Fig.1B, the viability of all four examined cell lines was modestly decreased by intervention with 5 μM *t*BHQ, but 10 μM of this chemical enabled these cell viability to return closely to their basal levels. Then, a relatively stable viability of *Nrf1α^−/−^* cells was maintained between 10-60 μM of *t*BHQ, followed by a gradual decrease to 80% of its viability until its concentration increased to 100 μM (Fig. 1B). By contrast, a narrow window of stable *WT* cell viability was defined by 10-20 μM *t*BHQ, followed by a fairly sloping downhill to 70% viability of the cells treated with 100 μM *t*BHQ, while other two close smoothly growth curves emerged from 10 to 80 μM *t*BHQ treatments of both *Nrf2^−/−ΔTA^* and *caNrf2^ΔN^* cell lines, before their viability decreased to 80% and 70%, respectively, upon treatment of 100 μM *t*BHQ (Fig. 1B).

Based on the dose-dependent effects, 50 μM *t*BHQ was selected for intervention of the above-described four cell lines to assess distinct time-dependent growth courses (Fig. 1C). The results showed that the viability of all four cell lines decreased to different extents of between 90% and 75% by *t*BHQ intervention for 1 h. Of note, the continuous treatment enabled the viability of *WT* and *Nrf2^−/−ΔTA^* cell lines to smoothly decrease to 85%-80% or 75%-75% from 2 h or 4 h to 24 h, respectively (Fig. 1C). By sharp contrast, the viability of *Nrf1α^−/−^ and caNrf2^ΔN^* cell lines appeared to elevate respectively to 100% or 90% in a modest ‘bounce-back’ response to *t*BHQ-continued treatment from 2 h to 4 h, and then both declined to 75% at 12 h of treatment. Thereafter, the viability of *Nrf1α^−/−^* cells continued to gradually reduce to 70% until 24 h of *t*BHQ treatment, whilst the viability of *caNrf2^ΔN^* cells was maintained to 75% from 12 h to 24 h treatments. As such, all cell viability reached a relatively stable level of them after 16 h of *t*-BHQ intervention. Therefore, the optimal concentration of *t*BHQ and its optimal time course were selected in the follow-up experiments to assess the cytoprotective roles of *Nrf1* and *Nrf2* against this chemical. For this end, we mainly investigated their expression differences between these four cell lines in responses to 50 μM *t*-BHQ intervention for 0, 1, 2, 4 and/or 16 h.

### 3.2 Short-term intervening effects of tBHQ on Nrf1, Nrf2 and AREs-driven genes in distinct genotypic cells

Herein, short-term effects of *t*BHQ intervention for 1-2 h on *Nrf1* and *Nrf2* were examined by western blotting (Fig. 1, D & E). The results showed that *t*BHQ treatment of *WT* cells caused modest increases in Nrf1-processed isoforms C/D, as well as *Nrf1^ΔN^* (Fig. 1D, *d1*). Such altered *Nrf1^ΔN^* also emerged in *Nrf1α^−/−^* cells, albeit it lacked A to D isoforms, implying it is not originated from the full-length *Nrf1α* processing. By contrast, a slight enhancement in the remnant *Nrf1α*-derived isoforms and *Nrf1^ΔN^* expression in *t*BHQ-treated *Nrf2^−/−ΔTA^* cells, but both their basal and *t*BHQ-stimulated levels were increased in *caNrf2^ΔN^* cells (Fig. 1D, *d1*). For *Nrf2*, its protein expression was more sensitive to *t*BHQ stimulation in *WT* cells, and also increased significantly after 1 h treatment (Fig. 1D, *d2*), when compared with those in the other three cell lines, which appeared to be largely insensitive to *t*BHQ, even although altered *Nrf2* expression levels were evidently enhanced in *Nrf1α^−/−^* and *caNrf2^ΔN^*, except *Nrf2^−/−ΔTA^*, cells lines, but with no obvious changes after *t*BHQ.

Both basal and *t*BHQ-stimulated expression levels of ARE-driven genes regulated by Nrf1 and/or Nrf2 were determined, next (Fig. 1, D & E). The results revealed that distinct expression levels of NQO1 (NAD(P)H:quinone oxidoreductase 1; Fig.1D, *d4*), GCLM (glutamate-cysteine ligase modifier subunit; Fig.1E, *e2*), GPX1 (glutathione peroxidase 1; Fig.1E, *e4*) and HO-1 (heme oxygenase 1, also called HMOX1; Fig.1E, *d4*) in *WT* cells were induced by *t*BHQ; this appeared to be accompanied by Nrf2 inducible enhancement. However, all these examined proteins, also including GCLC (glutamate-cysteine ligase catalytic subunit; Fig.1E, *e1*), GSR (glutathione-disulfide reductase, *e3*) and TALDO (transaldolase 1, *e5*), were largely unaffected by short-term *t*BHQ intervention of *Nrf1α^−/−^* cells, even though basal abundances of NQO1, GCLM and GPX1, amongst them aforementioned, were highly augmented as accompanied by hyper-expression of Nrf2. Similarly, constitutive active *caNrf2^ΔN^* also resulted in basal increases in GCLM, GPX1 and TALDO (Fig. 1E, *e2, e4, e5*) as accompanied by a basal decrease of NQO1, but all these examined protein levels were almost unaltered by *t*BHQ stimulation of *caNrf2^ΔN^* cells. Conversely, knockout of *Nrf2^−/−ΔTA^* only led to reduced basal levels of both HO-1 and GSR (Fig.1D, *d3* & Fig. 1E, *e3*), whilst *t*BHQ stimulation merely caused an inducible increase of TALDO alone in *Nrf2^−/−ΔTA^* cells (Fig. 1E, *e5*). Intriguingly, *t*BHQ-triggered *Nrf2^−/−ΔTA^* cells also gave rise to modest decreases of NQO1, GCLM, or GPX1 (Fig.1D, *d4* & Fig. 1E, *e2, e4*). Altogether, these indicate that Nrf1 and Nrf2 could make differential yet integral contributions to basal and *t*BHQ-inducible expression levels of these examined ARE-driven genes. For further insights into differential expression patterns of these antioxidant cytoprotective genes among different genotypic cell lines, the following experiments were performed by long-term stimulation of cells with *t*BHQ for 4 h to 24 h.

### 3.3 Long-term stimulating effects of tBHQ on Nrf1, Nrf2 and downstream targets in distinct genotypic cells

To give a proper understanding of long-term *t*BHQ-stimulated effects on *Nrf1, Nrf2* and downstream genes, their mRNA expression levels were determined by quantitative real-time PCR (Fig. 2). The results revealed an obvious increase of *Nrf1* mRNA expression after 12-h *t*BHQ stimulation of *WT* cells (Fig. 2A). By contrast, basal *Nrf1* mRNA expression was substantially abolished or diminished by knockout of *Nrf1α^−/−^* or *Nrf2^−/−ΔTA^*, respectively. Therefore, although Nrf2 was highly expressed in *Nrf1α^−/−^* cells (Figs. 1D & 2B), its remnant *Nrf1* was insensitive to *t*BHQ (Fig. 2A), whereas the residual *Nrf1* in *Nrf2^−/−ΔTA^* cells also hardly trigger a marginal response to this chemical. Conversely, *caNrf2^ΔN^* cells had given rise to a remarkable increase in basal *Nrf1* mRNA levels, but only a modest *t*BHQ-inducible increase of *Nrf1* expression was detected after 20-24 h stimulation of this cell lines (Fig. 2A). These results indicate that transcriptional expression of human *Nrf1* gene is monitored by Nrf2, as well by Nrf1 itself.

**Figure 2.**
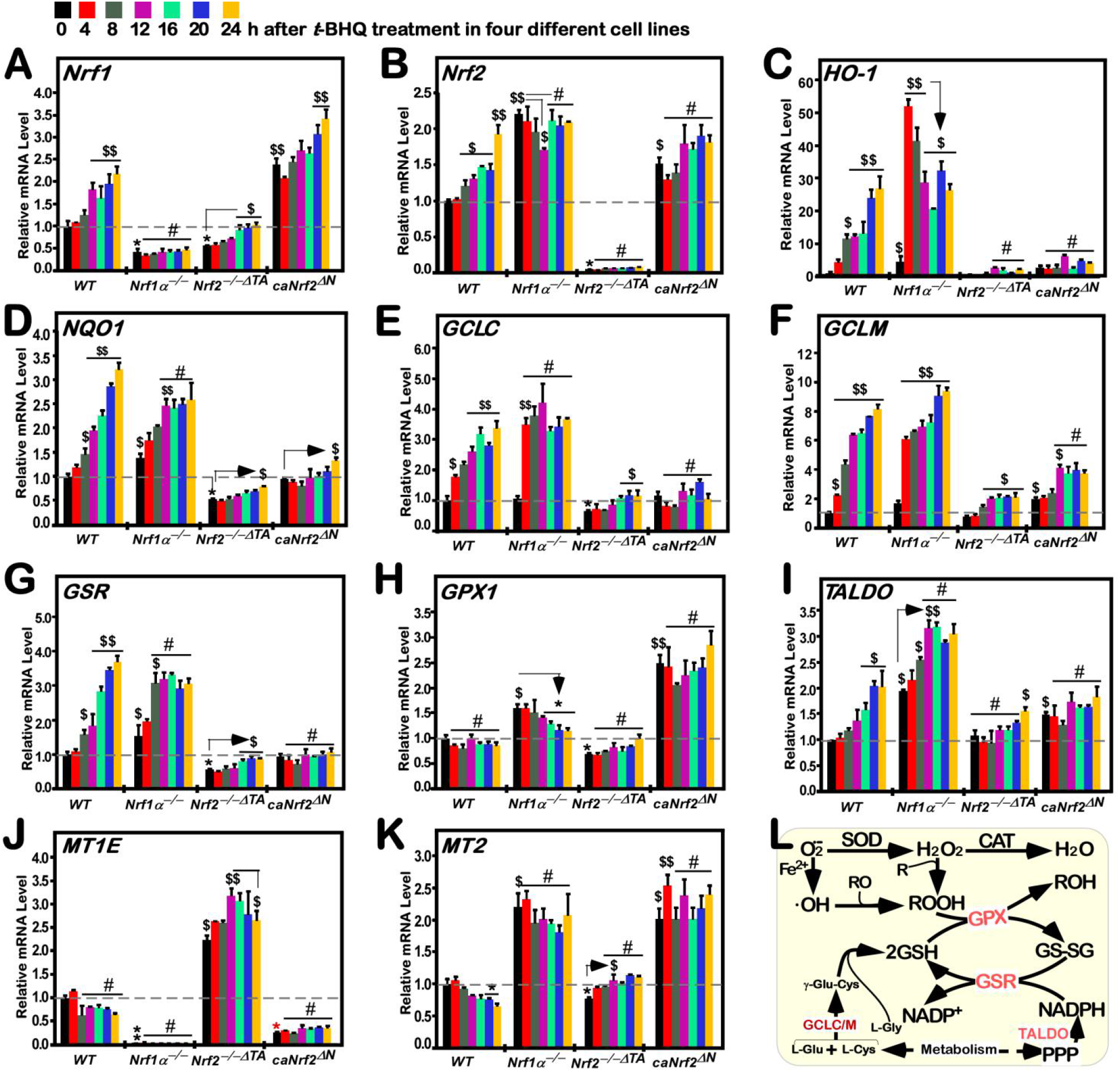
Time-dependent changes in the mRNA expression of distinctive responsive genes to tBHQ. Distinct genotypic cell lines of *WT*, *Nrf1α^−/−^, Nrf2^−/−ΔTA^* or *caNrf2^ΔN^* or were not treated with 50 μM *t*BHQ for 0 to 24 h, before both basal and *t*BHQ-inducible mRNA expression levels of all examined genes were determined by RT-qPCR. These genes included *Nrf1* **(A)**, *Nrf2* **(B)**, HO-1 **(C)**, NQO1 **(D)**, GCLC **(E)**, GCLM **(F)**, GSR **(G)**, GPX1 **(H)**, TALDO **(I)**, MT1E **(J)** and MT2 **(K)**. Then, statistic significances were calculated as described in “Materials and Methods”. Significant increases ($, *p*<0.05; $$, *p*<0.01), and significant decreases (\**P*<0.05; \*\**P*<0.01), were indicated, respectively, in addition to the non-significance (#). **(L)** A schematic representation of major ROS-scavenging enzymes and relevant redox signaling.

Treatment of *WT* cells with *t*BHQ caused a gradual modest induction of *Nrf2* mRNA expression levels from 8 h to16 h, followed by a marked peak of its induction at 24 h, of this chemical stimulation (Fig. 2B). Both basal and *t*BHQ-stimulated *Nrf2* expression levels were abolished by *Nrf2^−/−ΔTA^*. Interestingly, even though basal *Nrf2* mRNA expression was significantly augmented by *Nrf1α^−/−^* or *caNrf2^ΔN^*, its *t*BHQ-stimulated expression was unaffected or partially reduced in such two distinct genotypic cell lines (Fig. 2B). Collectively, these indicate that transcriptional expression of human *Nrf2* gene is bidirectionally regulated by itself and Nrf1, but upon stimulation by *t*BHQ, itself regulation by Nrf2 *per se* appears to be attributable to its N-terminal Keap1-binding Neh2 domain.

Besides Nrf1 and Nrf2, downstream target genes *HO-1* and *NQO1* were also induced by *t*BHQ of *WT* cells in a time-dependent manner (Fig. 2, C & D). Upon loss of Nrf1α-derived isoforms, significant increments in basal and *t*BHQ-stimulated mRNA expression levels of *HO-1* and *NQO1* were determined in *Nrf1α^−/−^* cells. The first sharp maximum peak of *HO-1* occurred at 4 h of induction by *t*BHQ, followed by a gradual decline to 16 h and then the second peak at 20 h treatment of *Nrf1α^−/−^* cells (Fig. 2C). By contrast, only a smooth increase of *NQO1* was obtained from 4 h to 12 h of *t*BHQ induction of *Nrf1α^−/−^* cells to a higher level, which was then maintained at such high level until 24 h (Fig. 2D). However, loss of *Nrf2^−/−ΔTA^* led to an evident diminishment or abolishment in basal and *t*BHQ-stimulated expression levels of *HO-1* and *NQO1* (Fig. 2, C & D), except for a marginal induction of *NQO1* by *t*BHQ at 24 h. Conversely, constitutive active *caNrf2^ΔN^* appeared to have no significant effects on both basal and *t*BHQ-stimulated expression of *HO-1* and *NQO1* (Fig. 2, C & D), albeit with a weak induction of *NQO1* by *t*BHQ at 24 h. Altogether, these data indicate that *HO-1* and *NQO1* serve as two representative targets of Nrf2, and both gene regulation by Nrf2 may also be monitored positively by its N-terminal Neh2 domain, aside from the potential negative regulation of Nrf2 and its targets by Nrf1.

### 3.4 Long-term stimulation of human antioxidant and detoxification genes by tBHQ in distinct genotypic cells

It is of crucial antioxidant and detoxification to be merited by glutathione (GSH)-conjugates in redox signaling cycles. The intracellular biosynthesis of GSH, as an important cellular antioxidant, is controlled by a key rate-limiting enzyme consisting of both GCLC and GCLM subunits. As shown in Fig. 2 (E & F), a time-dependent increment in the mRNA expression of *GCLC* and *GCLM* induced by *t*BHQ from 4 h to 24 h was determined in *WT* cells. By contrast, *Nrf1α^−/−^* cells gave rise to a rapid induction of *GCLC* mRNA expression by 4-h of *t*BHQ stimulation, to a maximum peak similar to that of *WT* cells, which was then maintained at such higher level until 24 h (Fig. 2E). However, no striking changes in both basal and *t*BHQ-stimulated *GCLC* expression were observed in *Nrf2^−/−ΔTA^* or *caNrf2^ΔN^* cell lines. Interestingly, further examinations revealed that basal and *t*BHQ-stimulated *GCLM* expression levels were substantially augmented in *Nrf1α^−/−^* cells (Fig. 2F), whilst *caNrf2^ΔN^* cells only gave rise to a relatively lower induction of *GCLM* by *t*BHQ, even though its basal expression was elevated at a similar level to that obtained from *Nrf1α^−/−^* cells. In addition, *Nrf2^−/−ΔTA^* cells still retained a considerable lower induction of *GCLM* by *t*BHQ (Fig. 2F). Taken together, these results demonstrate differential contributions of Nrf1 and Nrf2 to basal and inducible regulation of both *GCLC* and *GCLM* genes controlling GSH biosynthesis in distinct genotypic cells.

As a central enzyme of the intracellular antioxidant defense, GSR can reduce the oxidized glutathione disulfide (GSSG) to the sulfhydryl form (GSH). In such a thiol-based redox cycle, another key enzyme GPX1, belonging to the glutathione peroxidase family, can catalyze the glutathione to reduce hydrogen peroxide (H2O_2_) and other organic hydroperoxides, in order to detoxify the oxidants and hence protect the cells from oxidative damages. Thereby, we examine the intervening effects of *t*BHQ on induction of *GSR* and *GPX1* mRNA expression mediated by Nrf1 and/or Nrf2 in distinct genotypic cell lines. As anticipated, RT-qPCR results revealed that *GSR* mRNA levels were strikingly gradually up-regulated by *t*BHQ stimulation of *WT* cells from 8 h to 24 h (Fig. 2G), while expression of *GPX1* was unaffected by this chemical treatment (Fig. 2H). Upon knockout of *Nrf1α^−/−^*, basal mRNA levels of *GSR* and *GPX1* were markedly enhanced, but only modest induction of *GSR*, rather than GPX1, by *t*BHQ occurred from 4 h to 8 h and thereafter maintained at a maximum level that was lower than equivalent values measured from *t*BHQ-treated *WT* cells (Fig. 2,G & H). Such *t*BHQ-trigged induction of *GSR*, as well as its basal expression levels, was substantially attenuated or abolished by knockout of *Nrf2^−/−ΔTA^* (Fig. 2G). However, *GPX1* was smoothly down-regulated by *t*BHQ stimulation of *Nrf1α^−/−^* cells (Fig. 2H), although hyper-expression of Nrf2 was preserved in this knockout cell line (Fig. 2B), but this effect appeared to be completely prevented by *Nrf2^−/−ΔTA^* (Fig. 2H). Conversely, constitutive active *caNrf2^ΔN^* only gave rise to a remarkable increase in basal expression levels of *GPX1*, but not *GSR*, except that both genes were almost insensitive to *t*BHQ stimulation (Fig. 2, G & H).

Furthermore, TALDO is a key enzyme of the non-oxidative pentose phosphate pathway (PPP) providing ribose-5-phosphate for nucleic acid synthesis and NADPH for lipid biosynthesis [42]. Notably, NADPH arising from this pathway enables glutathione to be maintained at a reduced state, thereby protecting sulfhydryl groups and cellular integrity from oxygen radicals. Herein, our results unraveled a stepwise inducible increase of *TALDO* mRNA expression levels from 4 h to 20 h of its maximum stimulation by *t*BHQ of *WT* cells, before being maintained until 24 h of this chemical stimulation (Fig. 2I). By contrast, *Nrf1α^−/−^* caused significant increments in basal and *t*BHQ-stimulated expression levels of *TALDO* from 4 h to 12 h of its rapidly inducible peak that was much higher than the values from *WT* control cells, which was largely retained to 24 h. Such induction of *TALDO* was almost abolished by *Nrf2^−/−ΔTA^* (except for a marginal induction of it by 24-h stimulation of *t*BHQ), and also suppressed by *caNrf2^ΔN^* (albeit its basal levels were augmented) (Fig. 2I). Collectively, these indicate that Nrf1 and Nrf2 exert differential yet integral roles in mediating the aforementioned antioxidant cytoprotective genes against *t*BHQ.

In addition, it is worth mentioning that metallothioneins (MT) cannot only maintain the metal homeostasis in *vivo*, but also serve as a redox buffer for ROS and other free radicals to play an essential role in the cytoprotection [43]. However, our examinations of *MT1E* and *MT2* unraveled that both genes were not merely insensitive to *t*BHQ, but were modestly down-regulated by this chemical intervention of *WT* cells (Fig. 2, J & K). Of note, basal mRNA expression of *MT1E*, along with its inhibitory effect of *t*BHQ were markedly diminished or completely abolished in *caNrf2^ΔN^* or *Nrf1α^−/−^* cells, respectively (Fig. 2J), but both cell lines gave rise to a remarkable increase of basal *MT2* expression, aside from that *t*BHQ-stimulated expression of *MT2* was elevated in *caNrf2^ΔN^* rather than *Nrf1α^−/−^* cells (Fig. 2K). By contrast, *Nrf2^−/−ΔTA^* cells gave rise to significant increases in basal and *t*BHQ-inducible mRNA expression profiles of *MT1E* from 4 h to 12 h of this chemical stimulation, prior to being maintained at a considerable higher levels until 24 h (Fig. 2J), whereas basal *MT2* expression level was modestly down-regulated by *Nrf2^−/−ΔTA^*, but with a marginal induction by *t*BHQ (Fig. 2K). Taken together, these suggest distinct contributions of Nrf1 and Nrf2 to transcriptional regulation of *MT1E* and *MT2*, respectively.

### 3.5 Distinct time-dependent effects of tBHQ on Nrf1, Nrf2 and target gene expression in different cell lines

As shown in Fig. 3 (A, *a1*), Nrf1α-derived isoforms A to D were obviously enhanced after 4 h of *t*BHQ treatment in *WT* cells and then maintained to their considerably higher extents between 8 h and 24 h, as illustrated graphically (Fig. 3A, *a6*). Similarly, the abundance of Nrf2 proteins was rapidly significantly augmented by *t*BHQ stimulation of *WT* cells from 1 h to 24 h (Figs. 1D, *d2* & 3A, *a2*). Although hyper-expressed Nrf2 was retained in *Nrf1α^−/−^* cells, its protein abundances were unaffected by *t*BHQ intervention of this Nrf1α-specific knockout cells (Fig, 2B, *b2, b6*), in which Nrf1α-derived isoforms were constitutively lacked, but the N-terminal portion-truncated Nrf1^ΔN^ abundances were promoted by *t*BHQ (Fig, 2B, *b1*). By contrast, *Nrf2^−/−ΔTA^* cells only yielded considerably weaker abundances of Nrf1α-derived isoforms A to D, but they still enabled to respond *t*BHQ in a biphasic manner, with the first peak at 4 h of this stimulation and the recurring second peak at 20 h (Fig. 2C, *c1 & c6*), whilst the remnant Nrf2^ΔTAD^ protein was evidently time-dependently inhibited by *t*BHQ intervention of *Nrf2^−/−ΔTA^* cells (Fig. 2C, *c2*). Conversely, *caNrf2^ΔN^* cells led to strikingly increased abundances of basal Nrf2^ΔN^ and Nrf1α-derived isoforms (Fig. 1D, *d2* & *d1*), and also their time-dependent inducible expression changes were herein determined in *t*BHQ-stimulated *caNrf2^ΔN^* cells (Fig. 3D, *d1, d2* & *d6*).

**Figure 3.**
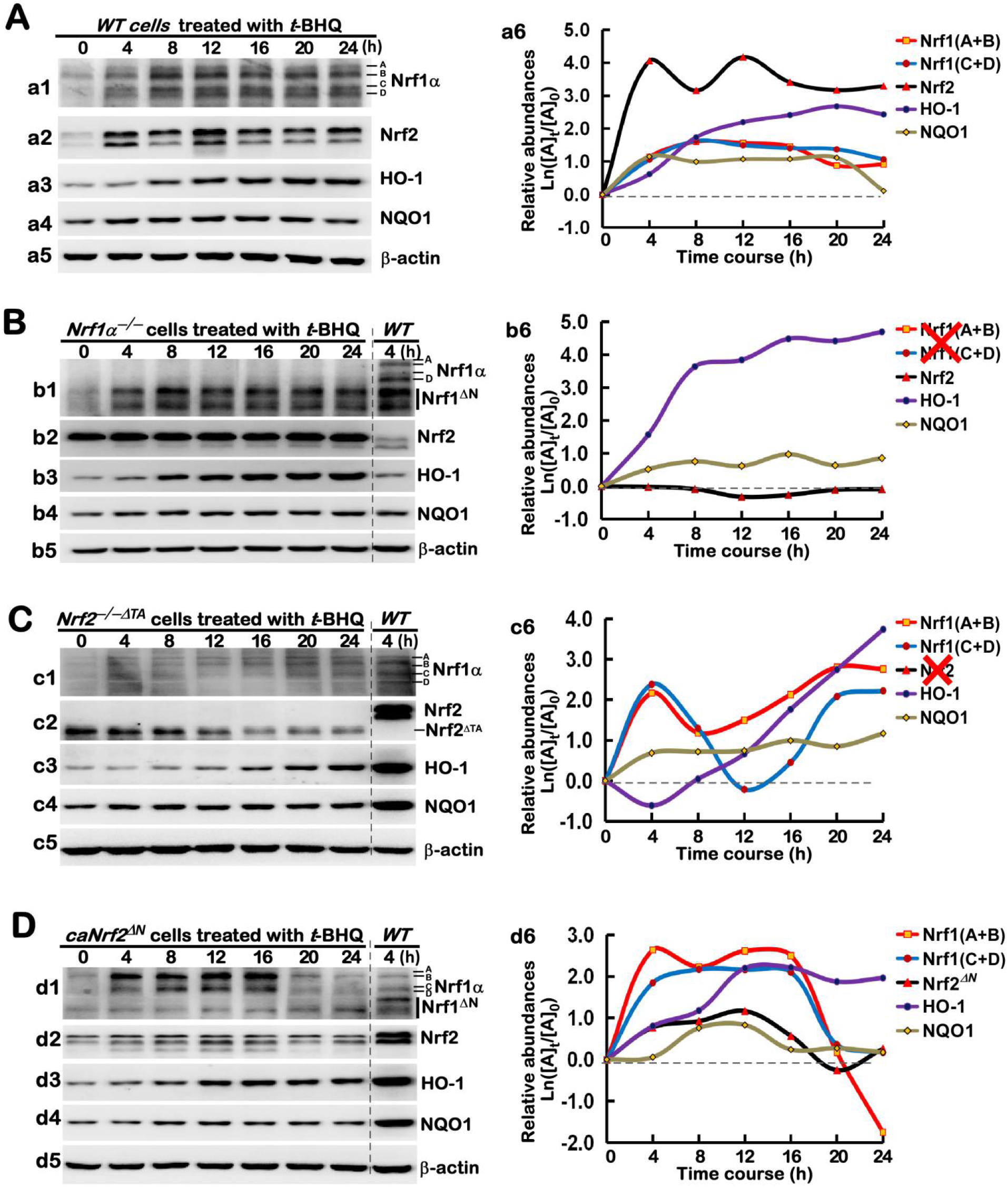
Long-term effects of *t*BHQ on protein abundances of Nrf1, Nrf2 and co-target genes. Experimental cells, *WT* **(A)**, *Nrf1α^−/−^* **(B)**, *Nrf2^−/−ΔTA^* **(C)** and *caNrf2^ΔN^* **(D)**, were or were not treated with 50 μM *t*BHQ for 0 to 24 h, before basal and *t*BHQ-inducible protein changes of Nrf1, Nrf2, HO-1 and NQO1 were determined by western blotting with their indicated antibodies, whilst β-actin served as a loading control. The intensity of those immunoblots representing different protein expression levels was also quantified by the Quantity One 4.5.2 software (Bio-rad, Hercules, CA, USA), and then shown graphically (in *right panels*). In addition, it should be noted that two big red crosses represent the constitutive losses of *Nrf1* or *Nrf2* (**b5** and **c5**), respectively.

Next, further examinations revealed that *t*BHQ-inducible expression levels of HO-1 in *WT* cells were gradually incremented, as its intervening time extended from 4 h to 20 h, to a considerably higher level, before being slightly declined (Fig. 3A, *a3* & *a6*), whilst NQO1-induced expression levels were rapidly triggered by 4-h of this stimulation to a certain extent, and then maintained until 24 h (Fig. 3A, *a4* & *a6*). Of great note, although basal abundance of HO-1 was substantially diminished by *Nrf2^−/−ΔTA^* (Fig. 1D, *d3*), it remained to be significantly induced by *t*BHQ from 8 h to 24 h of stimulation to a maximum extent (Fig. 3C, *c3 & C6*). In *Nrf1α^−/−^* cells, even though the hyper-expressed Nrf2 was insensitive to *t*BHQ, both HO-1 and NOQ1 were still rapidly induced by this chemical from 4 h to 8 h of stimulation and then maintained to their respectively higher extents until 24 h (Fig, 3B, *b2-b4, b6*). Rather, only a marginal induction of *caNrf2^ΔN^* by *t*BHQ was observed, but this was accompanied by significant induction of HO-1 and NQO1, as well as Nrf1α-derived proteins, in their distinct time-dependent courses (Fig. 3D, *d1-d6*). Altogether these collective results imply that such two inter-regulatory factors of Nrf1 and Nrf2 could mediate differential yet integral responses to *t*BHQ intervention of different genotypic cell lines.

### 3.6 Different time-dependent effects of tBHQ on antioxidant cytoprotective gene expression in distinct cell lines

As shown in Fig. 4A, *t*BHQ stimulation of *WT* cells caused significant time-dependent induction of GCLC, GCLM, GSR, GPX1, and TALDO (*a1* to *a7*). Amongst them, GCLC was relatively slowly induced after 8 h of *t*BHQ stimulation and then gradually incremented to a maximum inducible extent at 20 h of this chemical treatment, before being slightly declined (*a1*, *a7*). By contrast, GCLM, GSR and TALDO was rapidly induced within 4 h of stimulation by *t*BHQ and then presented in stepwise ascend from 8 h to 24 h of this treatment (*a2*, *a3*, *a5 & a7*), whilst GPX1 induction by *t*BHQ appeared to rise and fall in wave modes (*a4, a7*). Such differences in these examined enzymes may be attributable to distinct involvement of their upstream factors (e.g., Nrf1 and Nrf2) in response to *t*BHQ.

**Figure 4.**
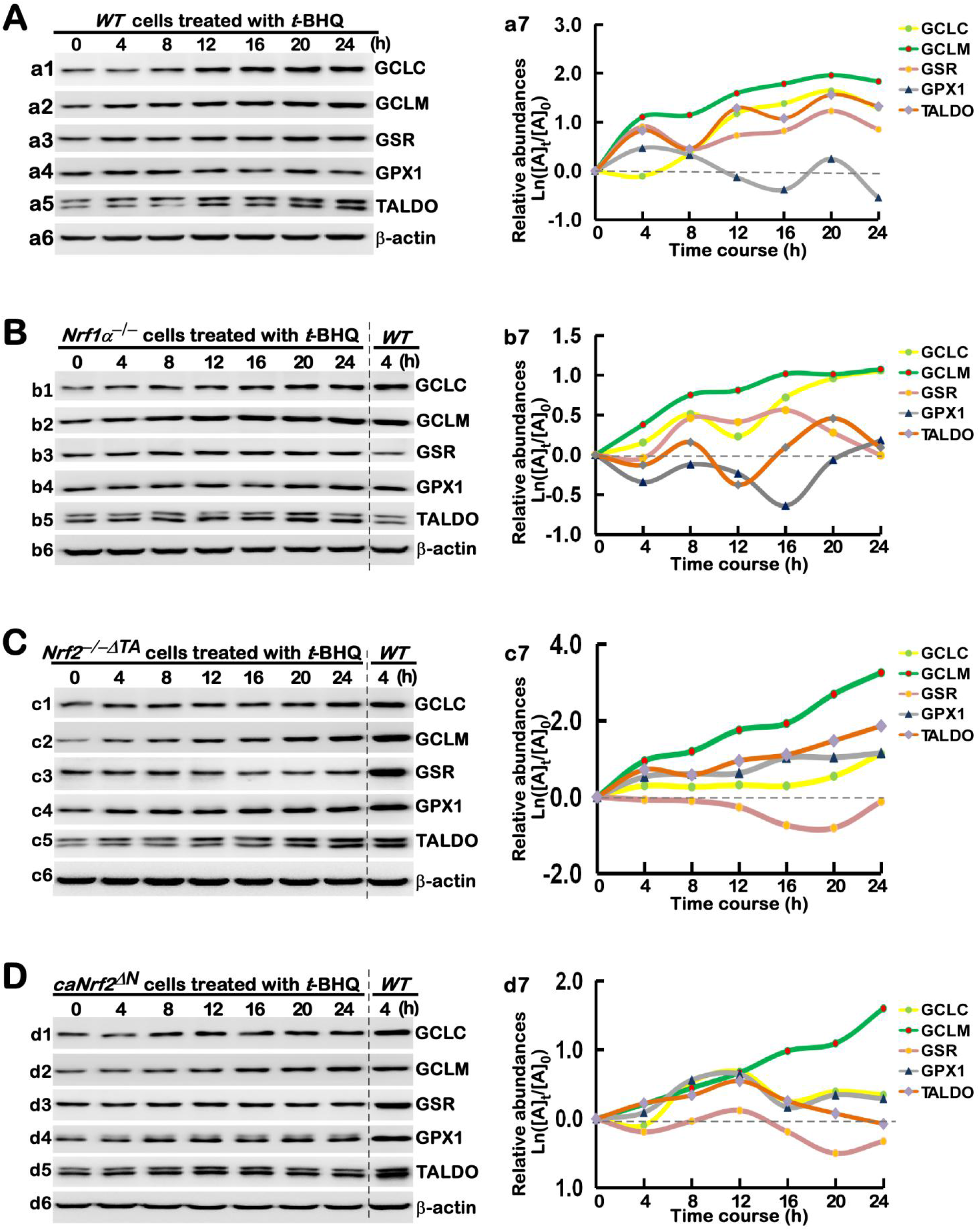
Long-term *t*BHQ-stimulated changes in the protein expression of antioxidant responsive genes. Different lines of *WT* **(A)**, *Nrf1α^−/−^* **(B)**, *Nrf2^−/−ΔTA^* **(C)** and *caNrf2^ΔN^* **(D)** were treated with 50 μM *t*BHQ or not for 0 to 24 h, before basal and stimulated abundances of those antioxidant cytoprotective proteins, e.g., GCLC, GCLM, GSR, GPX1 and TALDO, were determined by western blotting with the indicated antibodies. The intensity of relevant immunoblots representing different protein expression levels was quantified by the Quantity One 4.5.2 software, and then shown graphically.

Next, we determine distinct contributions of Nrf1 and Nrf2 to such alterations in *t*BHQ-stimulated abundances of antioxidant and detoxification enzymes. As revealed in *Nrf1α^−/−^* cells, GCLC and GCLM were successively induced by *t*BHQ from 4 h to 24 h of their maximum stimulation (Fig. 4B, *b1*, *b2 & b7*), while a lag induction of GSR occurred at 8 h of *t*BHQ stimulation, which was maintained to 16 h and then declined gradually to its basal levels (*b3*, & *b7*). As such, TALDO only displayed modest induction by *t*BHQ in a biphasic stepwise, with the first induction at 8 h and the second higher induction at 20 h before be declined nearly to its basal level (*b5 & b7*), except largely no induction of GPX1 in *Nrf1α^−/−^* cells (*b4 & b7*), although Nrf2 was hyper-expressed. However, *Nrf2^−/−ΔTA^* cells still gave rise to gradually-increased abundances of GCLC, GCLM, GPX1 and TALDO from 4 h to 24 h of *t*BHQ stimulation (Fig. 4C, *c1, c2 & c4-c7*), whereas GSR was slightly down-regulated by this chemical (*c3* & *c7*). By contrast, *caNrf2^ΔN^* only led to a significant increment in induction of GCLM by *t*BHQ from 4 h to 24 h (Fig. 4D, *d2* & *d7*), in addition to modest induction of GCLC, GPX1, TALDO, but not GSR, by this stimulation, which was maintained from 8 h to 12 h and then declined to relatively lower levels (*d1, d3-d7*).

### 3.7 Different antioxidant responses of four distinct genotypic cell line to tBHQ as a pro-oxidative stressor

The above experiments revealed there exists a synergistic effect of those antioxidant and detoxification genes regulated by Nrf1 and/or Nrf2. Just such synergistic effects can fully ensure the stable and effective function of this antioxidant cytoprotective system (to yield GSH and NADPH) to remove the excessive ROS produced from oxidative stressor, so that a certain redox homeostasis is being maintained to ensure the proper physiological operation of a heathy body. As a general term, ROS represents a set of all oxygen-containing reactive substances, including superoxide anion, hydrogen peroxide and relevant free radicals. To date, they remain to be hardly detected, owing to their characteristics of strong oxidative activity with such a short life to be rapidly scavenged and detoxified by antioxidants (i.e., GSH). As such, the intracellular redox state was herein measured directly by DCFH-DA, one of the most widely-used assays to evaluate the resulting oxidative damages, because it can react directly with ROS to give rise to an extremely sensitive, but impermeable, dichlofluorescein probe, as detected by flow cytometry [44]. As shown in Fig. 5A, a left shift of the dichlofluorescein image resulted from 16-h *t*BHQ intervention of *WT* cells (*a1*, also see Fig. S1), implying a relative decrease of intracellular ROS levels, when compared with the control image obtained from the vehicle treatment (at 0 h). Further examinations revealed that *Nrf1α^−/−^* or *Nrf2^−/−ΔTA^* gave rise to a significant increase in basal ROS levels under the vehicle-treated conditions, and also an evident left-shift of their *t*BHQ-intervening images to varying extents (Fig. 5A, *a2, a3* & 5B, and Fig. S1). This indicates that loss of stimulated by *t*BHQ to trigger antioxidant cytoprotective responses against the endogenic ROS. Intriguingly, almost no changes in both basal and *t*BHQ-stimulated dichlofluorescein images were determined in *caNrf2^ΔN^* cells, when compared to *WT* cells (Figs. 5A, *a3* & 5B, and Fig. S1), implying that the N-terminal Keap1-binding domain of Nrf2 is required for *t*BHQ-triggered antioxidant response.

**Figure 5.**
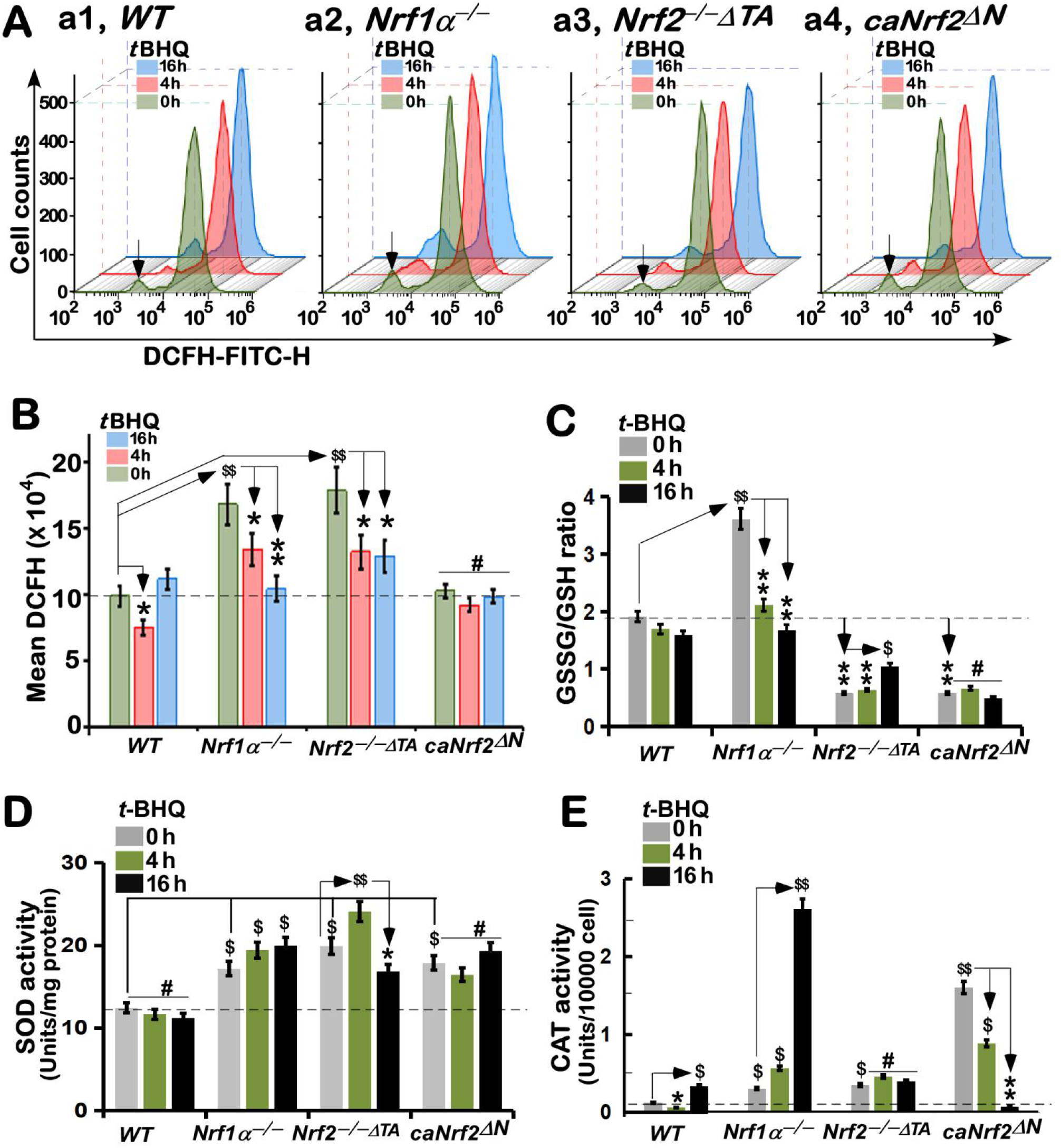
Altered levels of ROS, GSSG and GSH, along with ROS-scavenging activity of SOD and CAT, induced by *t*BHQ. **(A)** Experimental cells of *WT*, *Nrf1α^−/−^*, *Nrf2^−/−ΔTA^* and *caNrf2^ΔN^* were allowed for treatment with 50 μM *t*BHQ or not for different time periods (i.e. 0, 4, 16 h). Thereafter, the cells were subjected to a flow cytometry analysis of intracellular ROS by the DCFH-DA fluorescent intensity. The resulting data were further analyzed by FlowJo 7.6.1 software, as shown in the column charts **(B)**. Statistic significances were also calculated as described in “Materials and Methods”, with significant increases ($$, *p*<0.01), and significant decreases (\**P*<0.05; \*\**P*<0.01), respectively, in addition to the non-significance (#). **(C)** The intracellular GSH and GSSH levels, together with two ROS-scavenging activities of SOD **(D)** and CAT **(E)**, were measured according to the introduction of relevant kit manufacturers. The statistic significances were also determined as described above.

Further glutathione assays unraveled that the ratio of GSSG to GSH was marginally reduced by *t*BHQ stimulation of *WT* cells (Fig. 5C). By sharp contrast, *Nrf1α^−/−^* led to a remarkable increase in its basal GSSG to GSH ratio, but significant decreases of this ratio occurred after *t*BHQ stimulation. This indicates putative endogenous oxidative stress to yield the excessive GSSG, much more than GSH levels, in this Nrf2-hyperexpressed *Nrf1α^−/−^* cells, but the remaining antioxidant response in this cell line may be still triggered by *t*BHQ. Contrarily, *Nrf2^−/−ΔTA^* and *caNrf2^ΔN^* further caused substantial decreases in their basal GSSG to GSH ratio, although their stimulated ratios were less or not promoted by *t*BHQ, respectively (Fig. 5C). This implicates such two distinctive mutants enable to yield a certain amount of GSH in *Nrf2^−/−ΔTA^* and *caNrf2^ΔN^* cell lines, but could not allow for effective conversion of GSH into GSSG, even under *t*BHQ-stimulated conditions.

To gain insights into the initial scavengers of ROS, the activity of two key enzymes, superoxide dismutase (SOD) and catalase (CAT) was examined herein. As shown in Fig. 5D, significant increases in the basal activity of SOD were determined in *Nrf1α^−/−^*, *Nrf2^−/−ΔTA^* or *caNrf2^ΔN^* cell lines, and *t*BHQ-stimulated activity of SOD was further promoted only in *Nrf1α^−/−^, Nrf2^−/−ΔTA^*, but not *caNrf2^ΔN^*, cell lines. Of note, a longer term (16 h) treatment of *Nrf2^−/−ΔTA^* cells with *t*BHQ caused a substantial reduction of SOD activity to its basal level, which was, though, still higher than that measured from *WT* cells (Fig. 5D). However, *caNrf2^ΔN^* cells displayed no significant changes in *t*BHQ-inducible SOD activity, albeit its basal activity was much more than that of *WT* cells. Also, no obvious changes in the SOD activity were detected in *WT* cells that had or had not been treated by *t*BHQ. However, further examinations revealed that CAT activity was evidently stimulated by *t*BHQ in *WT* cells (Fig. 5E). By contrast, basal CAT activity was increased in both cell lines of *Nrf1α^−/−^* and *Nrf2^−/−ΔTA^*, but its *t*BHQ-stimulated activity was markedly elevated only in *Nrf1α^−/−^*, rather than *Nrf2^−/−ΔTA^*, cells (Fig. 5E). Furtherly, *t*BHQ stimulation of *caNrf2^ΔN^* cells caused a striking suppression or even complete abolishment of its inducible CAT activity at 4 h or 16 h of this treatment, respectively, although its basal activity was greatly substantially augmented. Such discrepant activities of SOD and CAT in between these cell lines, together with their differential expression results published previously [45], demonstrate to be attributable to distinctive yet cooperative contributions of Nrf1 and Nrf2 at regulating different target genes.

### 3.8 Distinct roles of Nrf1 and Nrf2 in different cell apoptosis induced by tBHQ as a pro-oxidative stressor

Further analysis by flow cytometry unraveled that only a few number of apoptotic cells were indeed examined in *WT* cells that had been intervened with *t*BHQ for 16 h (Fig. 6, A & B). By contrast, a remarkable augment in basal apoptosis of *Nrf1α^−/−^* cells reached to a much higher rate than that of the other cell lines, but its *t*BHQ-stimulated apoptosis was significantly decreased after intervention of *Nrf1α^−/−^* cells by this chemical for 4 h to 16 h (Fig. 6, A & B), to a similar level to that of *t*BHQ-treated *WT* cells. This phenomenon appeared to be almost consistent with the results of changing ROS levels as detected above (Fig. 5, A to C). These indicate that *t*BHQ induces antioxidant cytoprotective response against endogenous oxidative stress arising from *Nrf1α^−/−^* cells, in which putative hyper-expressed Nrf2 may be allowed for a certain extent to ameliorate potential oxidative damage and apoptosis caused by loss of Nrf1α, but could not fully compensate the constitutive loss of Nrf1’s function. Contrarily, no significant differences in basal apoptosis of *Nrf2^−/−ΔTA^* or *caNrf2^ΔN^* cell lines were observed when compared with that of *WT* cells (Fig. 6, A & B), but both mutants led to a modest or less increase in *t*BHQ-triggered apoptosis after intervention of *Nrf2^−/−ΔTA^* or *caNrf2^ΔN^* cells, respectively. This indicates that the sensitivity to *t*BHQ cytotoxicity may be weakened by *Nrf2* deficiency, allowing for the resistance of these two cell lines to a considerable extent of pro-oxidative stress. Overall, it could be concluded that both Nrf1 and Nrf2 play distinctive roles in mediating differential antioxidant cytoprotective responses against oxidative stress-induced apoptosis.

**Figure 6.**
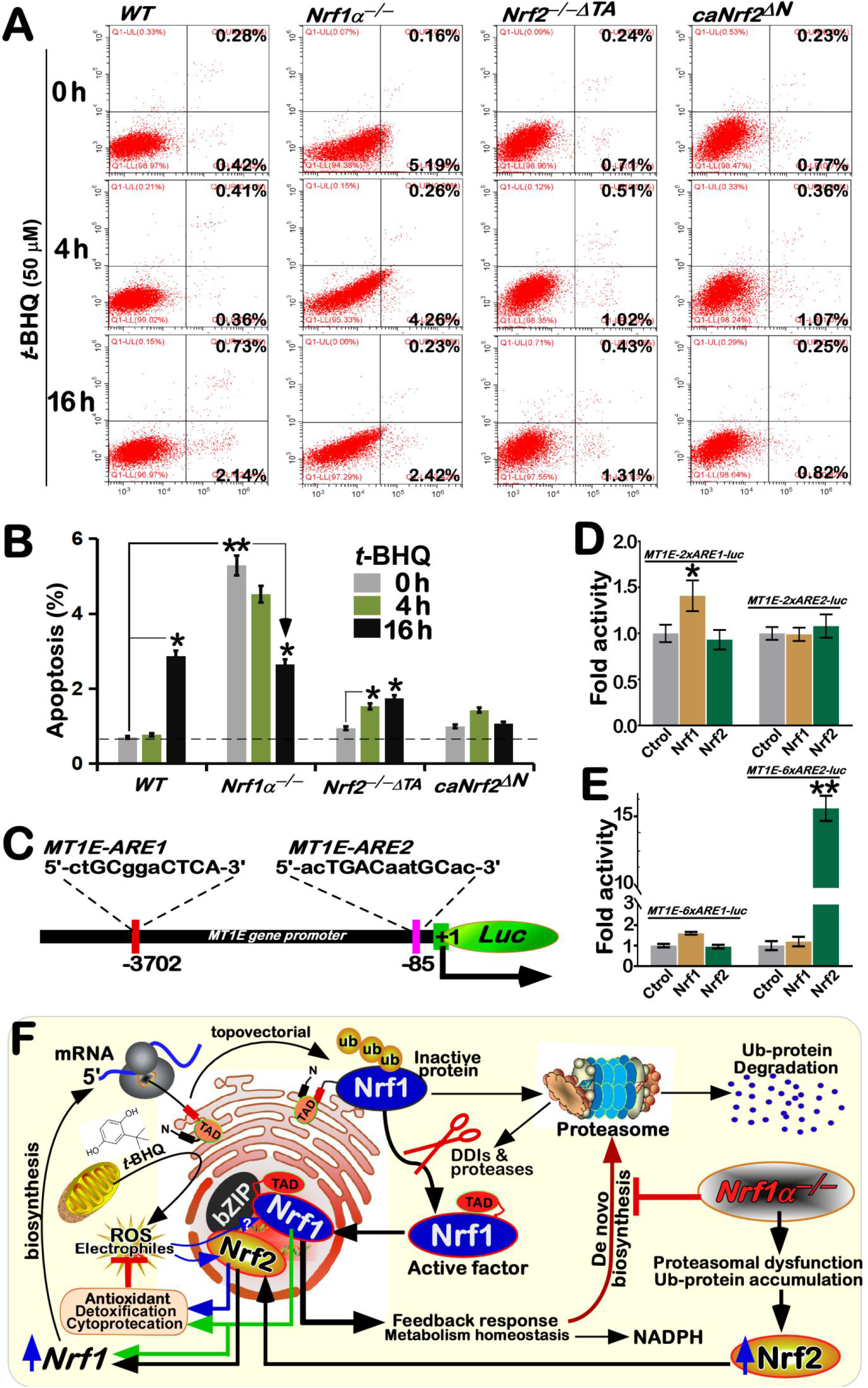
Nrf1 is more potent than Nrf2 at mediating the putative cytoprotective response to *t*-BHQ. **(A)** Distinct genotypic cell lines of *WT, Nrf1α^−/−^*, *Nrf2^−/−ΔTA^* and *caNrf2^ΔN^* were or were not treated with 50 μM *t*BHQ for different lengths of time. Subsequently, the cells were incubated with a binding buffer containing Annexin V-FITC and propidium iodide (PI) for 15 min, before being subjected to the flow cytometry analysis of apoptosis. **(B)** The final data were shown by the column charts. The statistic significances were also determined, with significant increases (\**P*<0.05; \*\**P*<0.01). **(C)** Two putative ARE sequences were shown in the location of the *MT1E* gene promotor region. **(D** & **E)** Distinct copies of those two ARE-sequences from the *MT1E* gene were allowed for driving relevant luciferase reporter genes. Each of ARE-driven reporters, together with expression constructs for Nrf1, Nrf2 or an empty plasmid, were transfected into *WT* cells for the indicated times. Then the cells were allowed for recovery from transfection, before the luciferase activity was measured, as shown graphically. The statistical determination was indicated with significant increases (\**P*<0.05; \*\**P*<0.01). **(F)** A proposed model is provided to give a better explanation of distinctive yet cooperative roles of Nrf1 and Nrf2 in synergistically regulating antioxidant cytoprotective genes. Of note, Nrf1 can act as a brake control for Nrf2’s functionality to be confined within a certain extent, albeit its transcriptional expression is also positively regulated by Nrf2.

This concluding notion is supported by further luciferase reporter assays (Fig. 6C), in which the reporter gene was driven by two different *ARE-*battery sequences existing in the promoter region of human *MT1E* (i.e., *MT1E-ARE1* and *MT1E-ARE2*). The results revealed that the transactivation activity of *MT1E-2xARE1-luc* was mediated by Nrf1 rather than Nrf2, but no changes in transcriptional expression of *MT1E-2xARE2-luc* were examined (Fig. 6D). However, a significant amplified activity of *MT1E-6xARE2-luc* was mediated by Nrf2 rather than Nrf1, even though the *MT1E-6xARE1-luc* was still modestly induced by Nrf1, but not Nrf2 (Fig. 6E). This difference in the activity of between *MT1E-2xARE2-luc* and *MT1E-6xARE2-luc* may be relevant to their contexts in the reporter gene constructs, but the detailed mechanism remains elusive.

## 4. Discussion

Since oxidative stress was initially formulated by Helmut Sies in 1985 and later redefined by Dean P. Jones in 2006 [46–48], an overwhelming number of publications by this conceptual term had been collected within at least 331,795 entries of the PubMed (https://pubmed.ncbi.nlm.nih.gov) until the middle of 2021. Such a continuously-heating topic on oxidative stress (and redox signaling) is open to arouse great concerns from researchers in distinct fields, but also is one of the most persistently-existing intractable problems to be addressed for health and disease, particularly in changing environmental conditions. Among its merits elicited by evoking biological stress responses, a steady-state redox balance is maintained within certain threshold ranges by cell respiration, aerobic metabolism and redox switches governing oxidative stress responses [46, 47]. However, the pitfalls of oxidative stress can also lead to indiscriminate use of this term as a global concept, but without a clear relation to redox chemistry, in each particular case. For the underlying molecular details, the major role in antioxidant defense is fulfilled by antioxidant enzymes, but not by small-molecule antioxidant compounds (e.g., *t*BHQ), in the cellular biochemical processes.

In all life forms, distinct cells can constantly generate a certain amount of ROS (and free radicals) during aerobic metabolism, so that its hormetic effects could be triggered in order to establish normal physiological cytoprotective mechanisms against oxidative damages. Of note, oxidative stress occurs in cells when ROS production overwhelms the natural antioxidant defenses and/or redox controls are disrupted [46, 47]. If oxidative stress is over-stimulated for a long term, the resulting damages lead to pathophysiological deterioration of many human chronic diseases, including cancer, diabetes, atherosclerosis, and neurodegenerative diseases [28, 49]. Thereby, to combat excessive production of ROS, all the cells have been evolutionarily armed with a series of innate powerful antioxidant defense systems. Among them is a set of essential antioxidant, detoxification and cytoprotective mechanisms governed by the CNC-bZIP family of transcription factors [12, 50, 51]. In mammalian cells, Nrf1 and Nrf2 are two principal CNC-bZIP factors to regulate target genes by specific ARE-binding sequences in their promoter regions. A large number of studies on Nrf2 revealed it functions as a master regulator of antioxidant response and relevant redox signaling [25]. Such versatile Nrf2 acts *de facto* as a promiscuous, but not essential player for ARE-binding to most of target genes [52], supporting the notion that Nrf2 is dispensable for normal growth and development [26]. As a matter of fact, Nrf1, rather than Nrf2, is a living fossil with its ancestral properties, because it shares a highly evolutionary conservativity with SKN-1, Cnc and Nach factors [51]. Like these ancient homologues [53, 54], Nrf1 is topologically dislocated across ER membranes and then processed to give rise to an N-terminally-truncated active factor, similar to Nrf2, before transactivating its target genes [55–57]. Hence, a unique conserved, indispensable role is fulfilled by Nrf1, but not by Nrf2, in maintaining the steady-state threshold of robust redox homeostasis.

The evidence has been provided in the present study, unraveling differential yet integral contributions of Nrf1 and Nrf2 to synergistic regulation of antioxidant cytoprotective genes at basal and *t*BHQ-inducible expression levels in wild-type (*WT*) cells. Specific knockout of *Nrf1α^−/−^* leads to severe endogenous oxidative stress as elicited by increased basal ROS levels; this is accompanied by increased ratios of GSSG to GSH and apoptosis. In *Nrf1α^−/−^* cells, Nrf2 was highly accumulated, but also cannot fully compensate loss of Nrf1α’s function in its basal cytoprotective response against endogenous oxidative stress, even though it had exerted partially inducible antioxidant response as the hormetic effect of *t*BHQ against apoptotic damages. By contrast, *Nrf2^−/−ΔTA^* cells were manifested by obvious oxidative stress, partially resulting from a substantial reduction of Nrf1 in basal and *t*BHQ-stimulated expression levels. However, the inactive *Nrf2^−/−ΔTA^* cells can be still triggered to mediate a potent antioxidant response to *t*BHQ, as deciphered by a significantly decreased ration of GSSG to GSH. Conversely, a remarkable increase of the Nrf1 expression was obtained from the constitutive active *caNrf2^ΔN^* cells, in which neither oxidative stress nor apoptotic damages had occurred, no matter if it was intervened with *t*BHQ. Thereby, distinct yet joint functions of Nrf1 and Nrf2 may be executed through their inter-regulatory effects on cognate genes against oxidative stress (Fig. 6F).

Differences in ROS-scavenging activities of SOD and CAT were determined in distinct genotypic cell lines. Basal activities of SOD and CAT were significantly increased by *Nrf1α^−/−^*, *Nrf2^−/−ΔTA^* or *caNrf2^ΔN^*, when compared to those of *WT* cells. *t*BHQ-inducible SOD activity were marginally elevated in *Nrf1α^−/−^, Nrf2^−/−ΔTA^*, but not *caNrf2^ΔN^* or *WT*, cell lines, as accompanied by an exceptional decrease of its activity by 16 h of this stimulation. The modest changes suggest that SOD activity may be monitored by other factors beyond Nrf1 and Nrf2. This notion is also supported by the previous RT-qPCR data [45]. Further evidence also revealed that *t*BHQ-stimulated CAT activity was markedly augmented in *Nrf1α^−/−^* cells (with hyper-active Nrf2 accumulation), and thereby completely abolished in *Nrf2^−/−ΔTA^* cells. However, basal increased CAT activity was substantially reduced by *t*BHQ intervention of *caNrf2^ΔN^* cells (with the enhanced expression of Nrf1), although this constitutive activator *per se* was unaffected by this chemical. These imply that CAT activity is regulated positively by Nrf2, and also monitored negatively by Nrf1, in particularly *t*BHQ-stimulated conditions. As such, it cannot also be ruled out that loss of the N-terminal Keap1-binding Neh2 domain from Nrf2 to yield *caNrf2^ΔN^* causes a negative effect on the *t*BHQ-stimulated CAT activity.

As a widely used Nrf2-activator, *t*BHQ can also trigger a certain activating effect on the expression of Nrf1 in *WT* cells, as well in *caNrf2^ΔN^* cells, but this effect is almost abolished by *Nrf1α^−/−^* or *Nrf2^−/−ΔTA^*. Conversely, induction of *Nrf2* expression by *t*BHQ occurred only in *WT* cells, but not in *Nrf1α^−/−^*, *Nrf2^−/−ΔTA^* or *caNrf2^ΔN^* cell lines, although its basal expression levels are significantly augmented in *Nrf1α^−/−^* or *caNrf2^ΔN^* cell lines. Together, with our previous data [31, 40], these indicate that Nrf1 has an ability to confine Nrf2 within a certain extent, albeit its transcriptional expression is positively regulated by Nrf2, which functions as a limited chameleon activator. Such inter-regulatory effects of Nrf1 and Nrf2 on antioxidant, detoxification and cytoprotective genes, such as *HO-1, NQO1, GCLC, GCLM, GSR, GPX1, TALDO, MT1E* and *MT2*, were further determined in distinct genotypic cell lines. As anticipated, the comprehensive experimental evidence has been provided herein, unraveling that *HO-1, NQO1, GCLC, GCLM, GSR* and *TALDO* were induced by *t*BHQ stimulation of *WT* cells, but *GPX1, MT1E* and *MT2* were not stimulated or even slightly suppressed by this chemical. By contrast, *Nrf1α^−/−^* cells were still allowed for *t*BHQ-increased expression of *HO-1, NQO1, GCLM* and *TALDO*, but with an exceptional decrease of *GPX1*, even though hyper-expressed Nrf2 was unaffected by *t*BHQ. This implies an additional involvement of other transcriptional factors beyond Nrf1 and Nrf2 in these gene response to *t*BHQ as a pro-oxidative stressor. Intriguingly, both basal and *t*BHQ-stimulated expression levels of *MT1E* were strikingly augmented in *Nrf2^−/−ΔTA^*, but not *Nrf1α^−/−^* or *caNrf2^ΔN^*, cell lines, whereas *MT2* was marginally induced by *t*BHQ in *caNrf2^ΔN^* cells. This finding implies that *MT1E*, but not *MT2*, may serve as an Nrf1-specific target gene, as further evidenced by its relevant reporter assays. Moreover, these was accompanied by a modest inducible enhancement of GLCM and Nrf1, whereas all other examined genes were, to lesser or no extents, stimulated by *t*BHQ in either *Nrf2^−/−ΔTA^* or *caNrf2^ΔN^* cell lines. These indicated that most of all other examined genes except *MT1E* are regulated primarily by Nrf2, but induction of its transactivation activity by *t*BHQ is also limited by its constitutive loss of the Keap1-binding Neh2 domain in the mutant *caNrf2^ΔN^* factor.

## 5. Concluding remarks

Dramatic research advances of the past 25 years, since a fascinating discovery by Itoh, *et al* [23], have witnessed an overwhelmingly preferential option for Nrf2 lonely focused in all relevant fields, whereas Nrf1 was almost totally ignored by all others except for a very few of groups. Such disproportionately-biased consequence has resulted in a general misunderstanding of Nrf2 as an only master hub of predominantly regulating antioxidant, detoxification and cytoprotective genes, regardless of the exiting facts that Nrf1, rather than Nrf2, is highly conserved with those more ancient SKN-1, Cnc and Nach factors [51], and can fulfill unique indispensable roles for cell homeostasis and organ integrity during life process. In the present study, together with our previous publications [31, 40], Nrf1 and Nrf2 are experimentally evidenced to elicit differential yet integral roles in mediating antioxidant cytoprotective responsive genes against pro-oxidative stress induced by *t*BHQ. The inter-regulatory effects of Nrf1 and Nrf2 on differential expression levels of antioxidant cytoprotective genes, e.g., *HO-1, NQO1, GCLC, GCLM, GSR, GPX1, TALDO, MT1E* and *MT2*, as well on the ROS-scavenging activities of SOD and CAT, were determined in depth. The collective results demonstrate that both Nrf1 and Nrf2 can make distinctive yet cooperative contributions to finely tuning basal and *t*BHQ-stimulated expression of target genes within the inter-regulatory networks. Overall, Nrf1 is allowed to act as a brake control for confining Nrf2’s functionality within a certain extent, albeit its transcriptional expression is positively regulated by Nrf2.

## Author contributions

Both Z.W. and Z. F. performed the experiments with help of K.L., collected all the relevant data, made draft of this manuscript with most figures and supplemental information. Y.Z. designed and supervised this study, analyzed all the data, helped to prepare all figures with cartoons, wrote and revised the paper.

## Acknowledgments

We are greatly thankful to Drs. Lu Qiu (Zhengzhou University, Henan, China) and Yonggang Ren (North Sichuan Medical College, Sichuan, China) for their involvement in establishing the indicated cell lines used in this study. We also thank to all other present and past members of Prof. Zhang’s laboratory (at Chongqing University, China) for giving critical discussion and invaluable help with this work. Notably, this study was funded by the National Natural Science Foundation of China (NSFC, with a key program 91429305 and other two projects 81872336 and 82073079) awarded to Prof. Yiguo Zhang.

